# Cell minor-axis length is a critical feature for breast cancer cell migration on straight, wavy, loop and grid microfibre patterns

**DOI:** 10.1101/2021.07.19.452928

**Authors:** Duo Zhang, Yaqi Sheng, Nicholas Piano, Theresa Jakuszeit, Edward Cozens, Lingqing Dong, Iek Man Lei, Wenyu Wang, Eugene Terentjev, Yan Yan Shery Huang

**Affiliations:** Department of Engineering, University of Cambridge, United Kingdom; Cavendish Laboratory, University of Cambridge, United Kingdom; The Affiliated Stomatologic Hospital, School of Medicine, Zhejiang University, P.R. China

**Keywords:** breast cancer, cell migration, extracellular matrix, topography, pattern

## Abstract

Cell migration plays an important role in physiological and pathological processes where the fibrillar morphology of extracellular matrice (ECM) could regulate the migration dynamics. To mimic the morphological characteristics of fibrillar matrix structures, low-voltage continuous electrospinning was adapted to construct straight, wavy, looped and gridded fibre patterns made of polystyrene (of fibre diameter ca. 3 μm). With microfibres deposited onto non-passivated surfaces, cells were permitted to explore their different shapes in response to the directly-adhered fibre, as well as to the neighbouring patterns. For all the patterns studied, analysing cellular migration dynamics of MDA-MB-231 (a highly migratory breast cancer cell line) demonstrated a switch in behaviour when the pattern features approach the upper limit of the cell minor axis. Our findings suggest that, although cells dynamically adjust their shapes in response to different fibrillar environments during migration, their ability to divert from an existing fibre track is limited by the size along the cell minor axis. We therefore conclude that the upper limit of cell minor axis might act as a guide for the design of microfibre patterns for different purposes of cell migration.

## 1. Introduction

Cell migration refers to the polarisation and locomotion of living cells. It can occur spontaneously and also be driven by a variety of biological, chemical and physical signals.^1–3^ Extensive efforts have been made to investigate the effects of physical signals on cell migration, including the role of surface topography and roughness that mimic characteristics of extracellular matrice (ECM).^4–11^ As the physical architecture and packing of native ECM can be complex, normally comprising multiple global patterns and local topographies, a reductionist experimental approach can be taken to provide mechanistic insights. For example, cell migration on linear tracks of fibronectin with widths narrower than the cell size, was shown to exhibit a 1D migratory behaviour, which more closely mimicked the cell migration in 3D fibrillar matrices than 2D migration on a plane.^12^ Based on the above, it was suggested that the 1D fibre topography could be used as an effective model for understanding cell migration dynamics in 3D fibrillar matrices.^12^ Linear tracks and straight fibre patterns have provided the foundation to recapitulate fibrillar morphology,^13,14^ thus it is of interest to evaluate how the features at a higher scale of fibre architecture, such as waviness, loops, and cross-junction grid patterns commonly found in ECM, could influence the cell migration dynamics.

To simplify typical ECM architectures into well-defined and repeatable patterns, this study adopted a low-voltage continuous electrospinning technique^15^ to design cell migration assays consisting of individual polystyrene microfibre tracks of straight, wavy, looped or gridded patterns. Microfibres were deposited onto non-passivated glass-coating substrates, thus cells had the freedom to explore different shapes in response to the well-defined fibril geometries. The combination of such fibre topography with ECM morpho-typical contours could potentially provide new insights into physically-guided cell migration behaviour, compared to established synthetic migration assays such as micro-patterned lanes^16^ and channels^17^. Here, we examined how different fibre patterns influence the migration dynamics of MDA-MB-231, a human breast cancer adenocarcinoma cell line characterised by elevated SNX27 (a protein marker of cell mobility and proliferation) expression, and the highest persistent 2D migration among all breast cancer cell lines^18–21^.

## 2. Results and Discussion

### Global fibre pattern determines cell migration direction and cell shape, but not the speed

Disease or age-associated extracellular matrix remodelling often leads to the creation of stiff collagen fibre bundles, which could impart biomechanical cues to cells with their rod-like cross-section (several microns in diameter) and nano-scaled surface topography.^22–24^ To create synthetic microfibres mimicking the physical attributes of individual stiff ECM fibre bundles, low-voltage continuous electrospinning patterning (LEP), developed in our previous work,^15,25^ was used to fabricate polystyrene fibres with required features. Made of polystyrene, such synthetic fibres could provide a chemically stable and robust framework to study cell migration. **Figure 1**(a) shows the basic fibre deposition process on glass slides. The diameters of fibres produced have an average diameter of 3.3 ± 1.1 μm **(Figure S2(a))**. In **Figure 1** (b), scanning electron images of single fibres show their circular cross-section and ‘nano-wrinkle’ surface features. In **Figure 1**(c), the contact Young modulus of the fibre determined by atomic force microscopy (AFM) was shown to be around 200 MPa, within the range of contact moduli of collagen fibrils (∼0.1–10 GPa) measured by AFM with different buffer solutions.^26,27^ Adjusting the stage collection speed and the application voltage, fibre contours could be made to mimic morphological characteristics of the extracellular matrix^28–31^, from straight fibres, to waves, and loops, can be fabricated (**Figure 1**(d)-(g)). To define the wavy fibre features, we fitted sinusoidal functions to the fibre contours, as will be described in later sections. For loop fibres, as the loops were approximated circular **(Figure S2 (e))**, the equivalent loop diameters were determined to be between 20 to 40 μm in our study. Finally, for grid fibres, the fibre intervals were varied between 0 to 120 μm in orthogonal directions.

**Figure 1.**
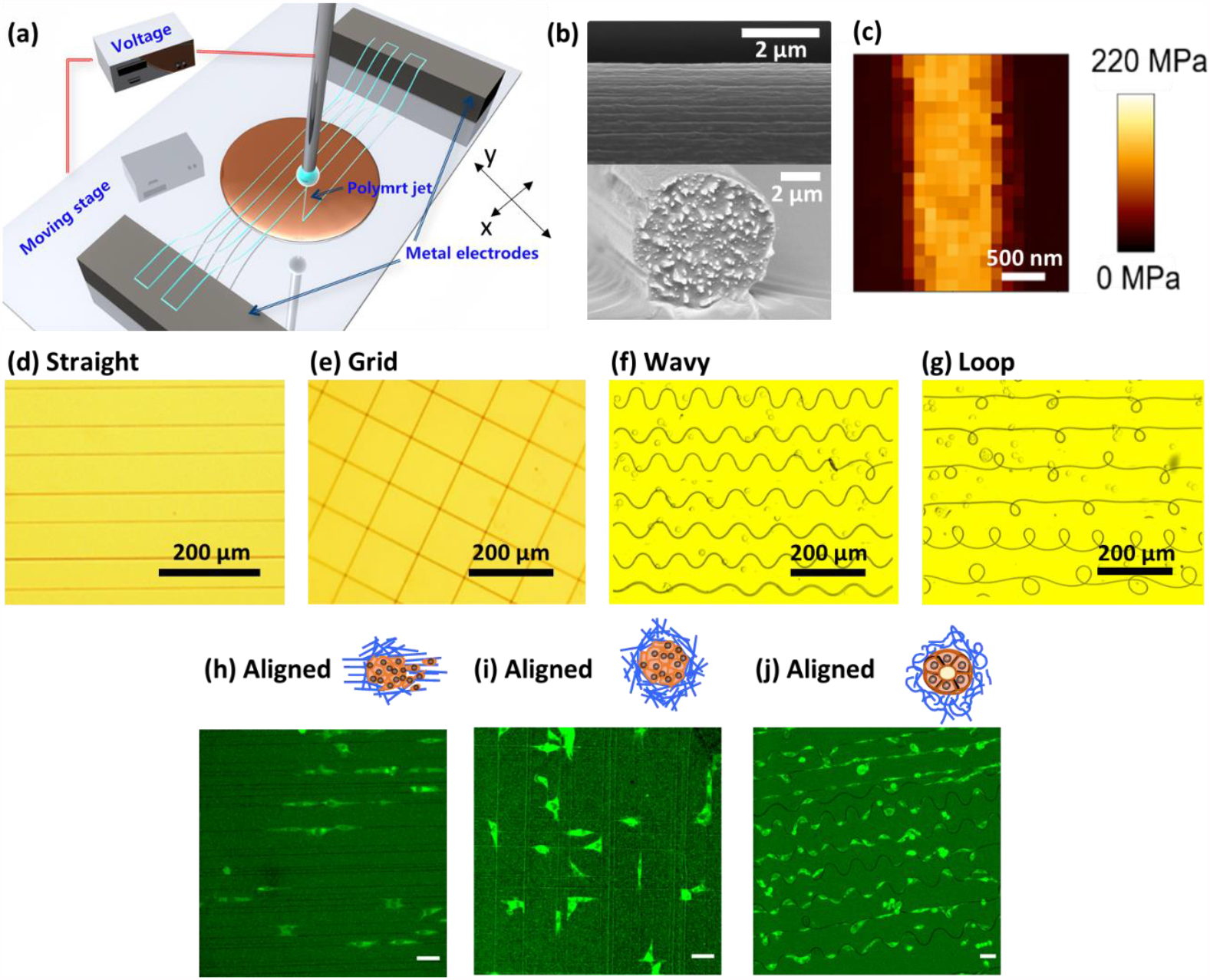
Overviews of the LEP fabricated polystyrene microfibres with different global patterns, as well as the cell-fibril migrations. (a) Schematic figure illustrates the equipment setup of LEP; (b) SEM images showing the nano-wrinkle features and the cross-sectional view of micron-sized fibres; (c) Apparent contact stiffness mapping measured by in situ AFM indentation on a polystyrene fibre; (d)-(g) Fabricated polystyrene fibres with straight aligned, grid, wavy and loop patterns; (h)-(j) Fluorescent microscopic images capturing single cells migrating along aligned, grid and curly fibre patterns, scale bar=50 μm.

Live cell imaging, with snapshot showed in **Figure 1**(h)-(j), confirmed that each individual fibre had the capacity to guide cell migration. Depending on the fibre pattern, cells adhering to individual fibres, or two parallel fibre tracks, perform guided movement along fibre axes. Cells on a network of orthogonal crossed fibres migrated along either horizontal or vertical directions. **Figure 2**(a) compares the average step speed (calculated from the distance travelled within 10.5 min of the imaging interval) for single cell migration on different fibre patterns. The average step speed ranges between 0.2-0.5 μm/min on the four typical global fibre patterns, with no significant difference found for single cell migration on grid, wavy and loop fibres, compared to that on straight aligned fibres. With the finding above, we suggest that the global fibre pattern affect the direction of cell movement, but not the individual cell step speed. Cell migration speed could be controlled by the fibre local topographies such as fibre dimension/ diameter and surface textures^32–35^, which were kept constant in our experiments. Next, analysing single cell morphology in **Figure 2**(b) revealed that as fibre patterns change from open linear to enclosed patterns, the cell aspect ratio increased to a maximum of 0.8 (approaching towards 1). Thus, the average shape of a cell is strongly influenced by the fibre pattern, which might have further implications for fibre-guided cell migration. In the following sections, we will analyse cell migration in each of the fibre pattern case in detail, and evaluate how the associated migration behaviour is correlated to the size and shape of the cells.

**Figure 2.**
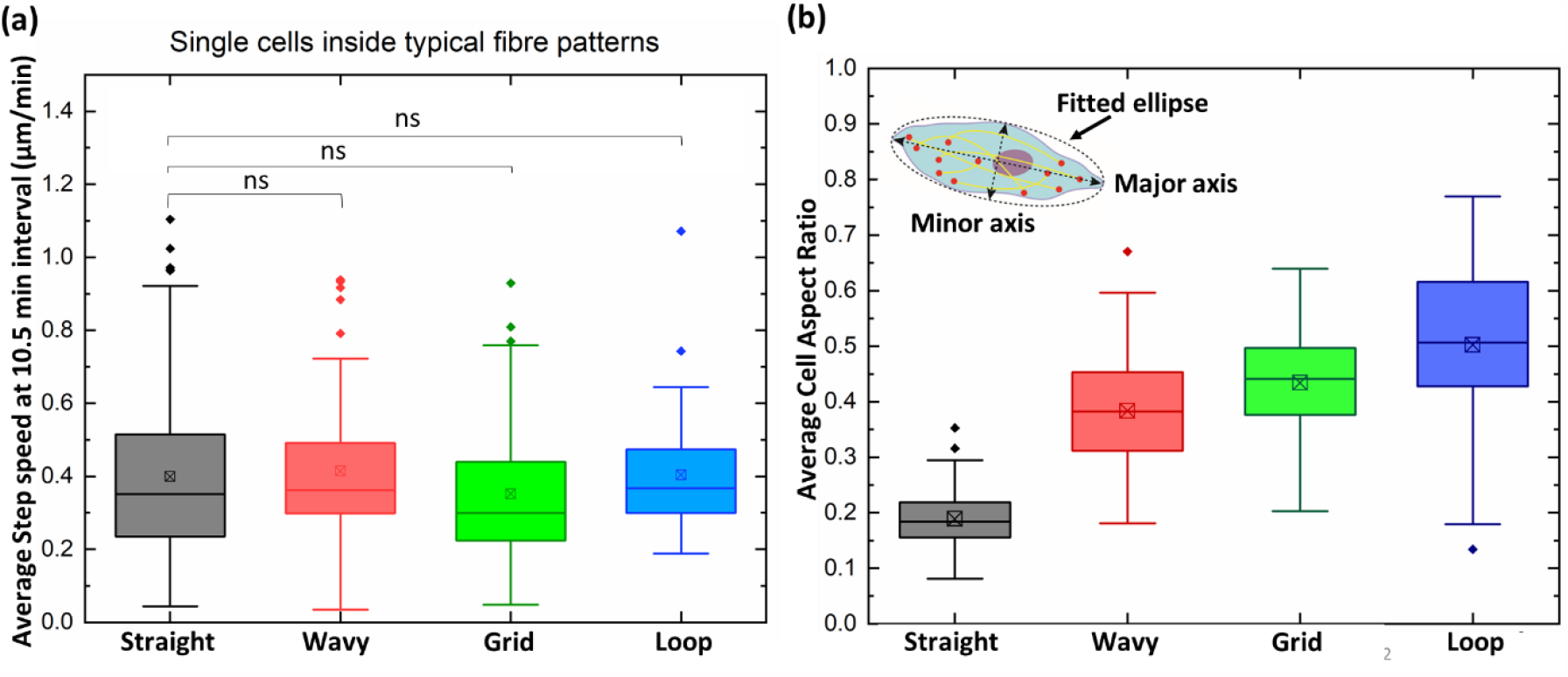
Cross comparison for single cells’ migration features along different fibre patterns. (a) Cross comparison of single cell step speed when migrating along different fibre patterns, step interval = 10.5min; (b) Cross comparison of single cell aspect ratio when migrating along different fibre patterns; with aspect ratio = minor axis length / major axis length. Number of single cells studied: n= 62 cells on straight fibres; n= 76 cells on wavy fibres; n= 98 cells on grid fibres; n= 39 cells on loop fibres.

### Migration dynamics along straight, parallel fibre patterns

For the straight fibres, topography guidance provided by the polystyrene tracks efficiently drove the cells to move along the fibre direction, as shown by **Figure 3**(a). During single cell migration, their major axis was aligned with the fibre axis. In some cases, a cell could stretch its body sideways and attempt to connect to multiple fibres during migration (bridging between parallel fibres); while in other cases, a cell remained solely on its original fibre track throughout the imaging period (see **Figure 3**(b)-(c)). Categorising these two scenarios of migration behaviour in **Figure 3**(d) suggests that the spacing between parallel fibres (i.e. fibre interval) is a determining parameter. When the fibre interval was smaller than 30 μm, there was no preferred choice for the cell behaviour (either remain on a single fibre or bridging to neighbouring fibres). However, when the fibre interval was increased to over 35 μm, cell bridging was no longer observed, and cells remained on their original linear fibre. **Figure 3**(d) shows that cell minor axis length all had values below 18 μm. Thus, we suggest that the adjacent fibre interval is the characteristic pattern size of the parallel fibres, which determines whether a cell would be diverted from its current fibre migration path. The length of cell minor axis could be used as a measure of their ability to stretch their body orthogonal to the fibre direction, to bridge an adjacent fibre for migration.

**Figure 3.**
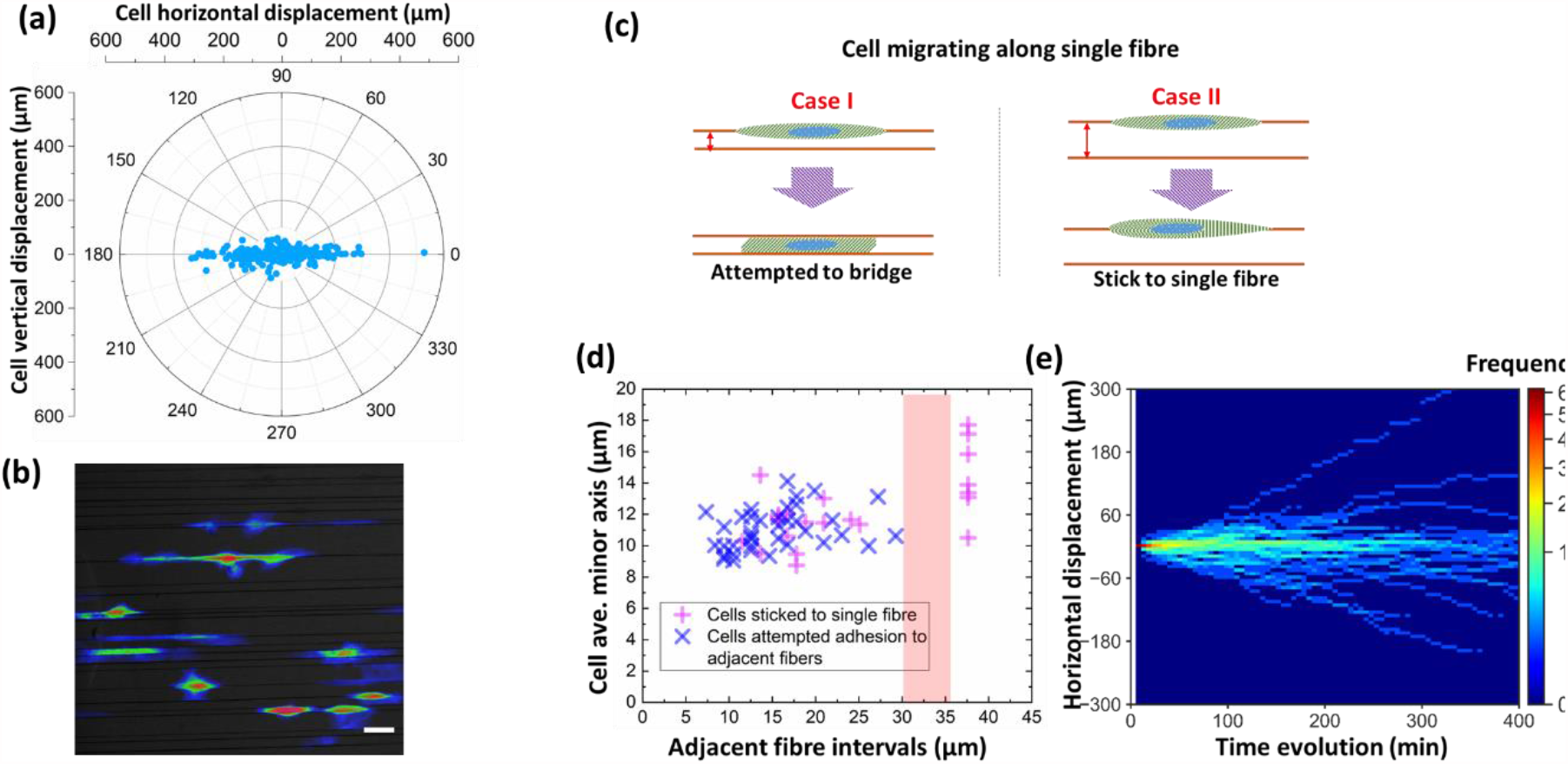
Analysis of cell migrations along aligned polystyrene microfibres. (a) The polar scattered plot demonstrating overall cells’ ultimate displacements on the aligned fibre patterns over 12-hour time evolution, with n=389 cells analysed in total, scale bar=50 μm; (b) A cumulative probability map demonstrating single cells migrating along straight aligned fibres over the12-hour imaging period; (c) Schematic diagram showing a single cell would either attempt to hold two parallel fibres (bridge) or stick to the original single fibre, depending on the adjacent fibre interval; (d) Adjacent fibre interval is considered as feature length, and the scattered plot shows the 30-35 μm range fibre interval is a threshold to differentiate single cell behaviours; (e) the 400-min time-accumulated displacement distribution for all single cells when they were attached to single fibres, with n=62 single cells analysed in total.

To evaluate the migration statistics for the single fibre migration case, cell displacement along the fibre direction was calculated as 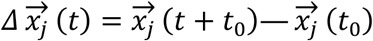 where 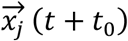 denotes the position vector (in our 1D system we only consider motion in one axis) of the j^th^ cell at time t, and t_0_ represents the corresponding original starting time. The displacement vs. time plot for all the cells tracked for the straight fibre patterns, is shown in **Figure 3**(e). The ensemble-averaged displacements of cells derived from all cell trajectories were centred approximately at zero-displacement, supporting the equal likeliness for the cells to migrate towards either end of the fibre.

### Migration dynamics along grid patterns

Looking over the 12-hour live-cell imaging period, an interesting observation is that most cells are restricted to their local grid blocks, which is shown in the fluorescent microscopic image as well as the probability map in **Figure 4**(a). The possible reason is that within the time of measurement, cells would inevitably come across orthogonal fibre joints, which appear as a local convex landscape to the cells. When a cell comes across an orthogonal fibre joint, its protrusions are able to sense orthogonal fibres, and the cell reacts accordingly to adapt to different fibril topography. The ultimate displacement of cells during the entire experimental session was calculated and displayed in **Figure 4**(b) (n=282 cells). Compared to cells on straight aligned fibres, this displacement distribution shows that grid joints provide an extra degree of freedom to cell migration.

**Figure 4.**
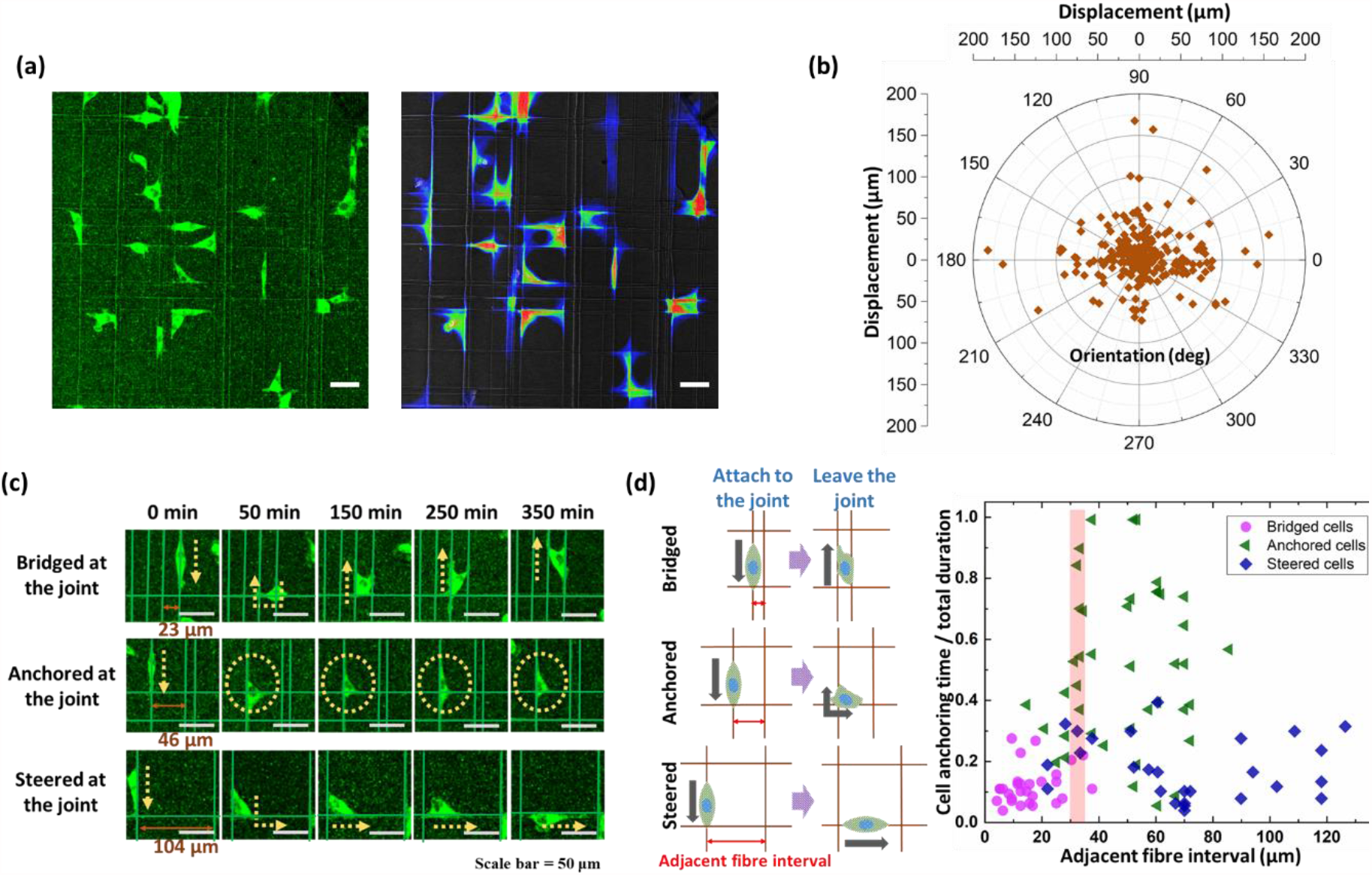
Analysis of single cell migration along grid polystyrene microfibre patterns. (a) Fluorescent microscopic image capturing cells migrating on fibre grids, and the cumulative probability map over the 12-hour imaging period, scale bar=50 μm; (b) The polar scattered plot demonstrating cells’ ultimate displacements on the grid fibre patterns over 12-hour time evolution, with n=282 cells analysed in total; (c) Set of diagrams demonstrating three typical scenarios of cell movements when migrating towards the grid fibre joints, scale bar=50 μm; (d) Schematics showing three typical scenarios for a single cell to interact with the fibre joints, as well as the correlation between adjacent fibre intervals and cells’ normalized residence. The threshold fibre interval (feature length) range is around 30-35 µm.

Next, the single cell migration was studied. **Figure 4**(c) illustrates fluorescent microscopic images of three types of cell behaviours that were observed when interacting with orthogonal fibre junctions: (i) bridged at joints, where cells would bridge to the adjacent parallel fibre track and then continue migrating along the two parallel tracks simultaneously; (ii) anchored at joints, where cells would remain anchored at these joints; and (iii) steered at joints, where other cells would be redirected along one of the orthogonal fibre tracks. To characterise the migration behaviour, we define the ‘cell anchoring time’, as the total time that a cell is anchored at a fibre junction, from the initial point the cell body first touches the joint, until the moment that all parts of the cell body have lost contact with the joint. When the normalisation of this parameter (cell anchoring time/total imaging time) is plotted against the adjacent fibre distance in **Figure 4**(d), the different behaviour patterns fall into distinct categories. When the adjacent fibre distance is less than 30-35 µm, a mixture of the three migration patterns was observed, though cell bridging dominates at narrow fibre spacing. As adjacent fibre distance becomes greater than 35 µm, cell bridging can no longer take place. For adjacent fibre distance above 70 µm, cells would then be redirected at these junctions to an orthogonal fibre track for continued migration.

### Migration dynamics along wavy and loop fibre patterns

ECM fibrils often have curly geometries, with features similar to cell sizes, and such geometries can be categorised into two major groups: wavy and closed-loop patterns. For wavy fibre patterns, the curve can be described by a sinusoidal function of y=A*sin(2πx/λ), with A being the amplitude and λ the wavelength. Focusing on single cell-fibre interaction, the migration along a wavy fibre segment is illustrated in **Figure 5**(a), where cells were observed to maintain contact with the fibre throughout the migration. A cell entering a wavy curve is defined to be the moment when the cell body firstly touch a trough/crest (where dy/dx = 0); and a cell leaving the wavy curve is defined as the moment when the cell body has fully passed over the curve trough/crest. Cell residence time represents the total time that a cell spent passing across a wavy fibre curve. **Figure 5**(b) displays three typical cases of microscopic images, showing different sizes of wavy fibre curves in bright field, and correspondent cell shapes in fluorescent channel. Probability map is used to illustrate the spatial distribution and activity of the cells over time. Live-cell images were captured at intervals of 10 min and overlapped into a 2D projection image, such that each image pixel numerically represents cell occupancy probability, with a value between 0 and 1. Cell residence times are also displayed alongside the probability maps, highlighting that as the waveform amplitude increase, coupled with increased curliness, cell residence seemed to be prolonged.

**Figure 5.**
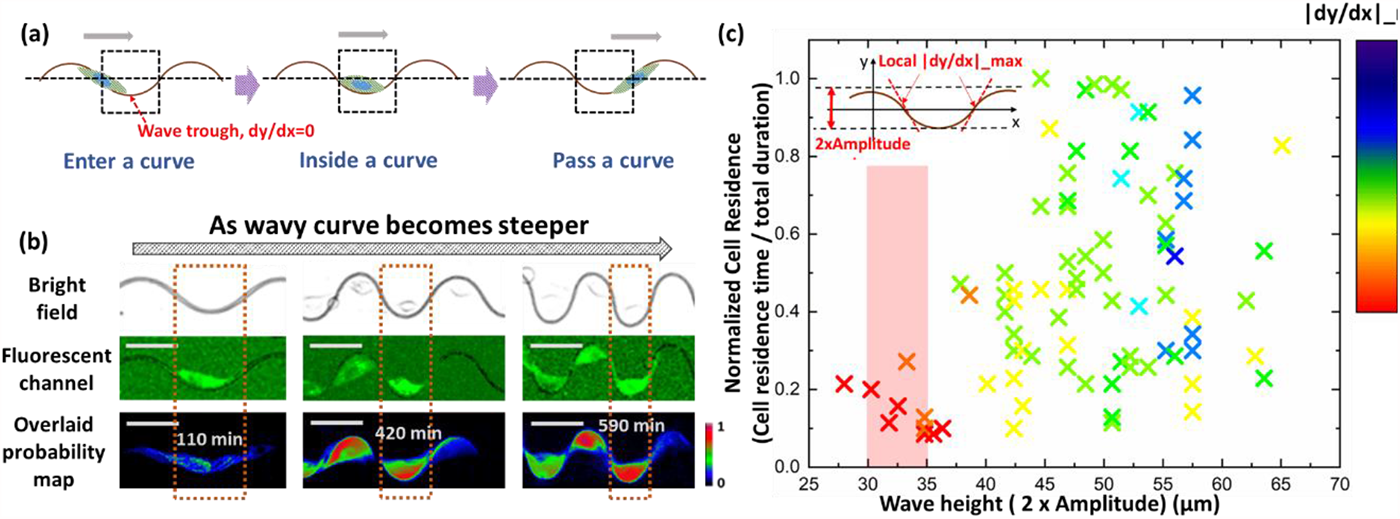
Analysis of single cell migrations along wavy polystyrene microfibre patterns. (a)Schematic diagrams illustrating different stages of a cell migrating across a wavy curve; (b)Cell behaviours differ as wavy curve become steeper, demonstrated in microscopic images in bright field, green fluorescent channel and time-accumulated probability maps, scale bar=50 μm; (c) Scattered distribution shows the wavy features (wave height and|dy/dx|_max) influence cells’ normalized residence. The 30-35 μm range in ‘wave height’ acts as a threshold that differentiates cell residence. ‘Wave height’ is defined as the difference between the elevations of a crest and a neighbouring trough (also twice the amplitude of the fitted sine wave).

To quantify the curliness of wavy fibres, ‘curve steepness’ was characterised as the maximum derivative (|dy/dx|_max) for each of the sinusoidal segment. Thus **Figure 5**(c) shows the effect of wavy fibre contour, characterised by wave height (2×A) and steepness (|dy/dx|_max) on cell residence. When the wave height increases beyond ∼30-35 µm, along with the increased curve steepness, the normalised cell residence times increased rapidly. In some cases, the ‘cell residence time/total time duration’ ratio approached 1, indicating that these cells were trapped inside the curve throughout the entire imaging period. It should be noted that as the curve steepness is increased, there are still cases of cells having lower residence time. This is because cells were observed to enter fibre curves at different times during the imaging session, thus low residence values were recorded for cells entering towards the end of the imaging session.

The close-loop patterns demonstrated an efficient ‘cell trapping’ ability when the loop diameters are close to the cell size. **Figure 6**(a) shows stages of cell transiting through a loop pattern, where we define in this case the cell residence time as the total time elapsed for a cell staying inside a loop. **Figure 6**(b) displays example microscopic images, showing different sizes of fibre loops in bright field, and correspondent cells in fluorescent channel. Accords with the schematic illustration back in **Figure 6**(a), for small loops with diameters of around 20 μm, cells tended to occupy the entire loop space. In contrast, for large loops, with diameters greater than 35 μm, cells could only adhere to a portion of the loop boundary, and would accordingly only occupy smaller portions of the loop. For medium size loops with diameters ranging between 20 to 30 μm, cells would still be able to accommodate the majority of the loop space; In addition, there tended to be different ways in which they would adhere to the loop boundary. **Figure 6**(c) presents the correlation between the normalized cell residence and the loop diameter. As the loop diameter increases from 20 to 35 μm, the cell residence gradually shifted up to around 70% of the time.

**Figure 6.**
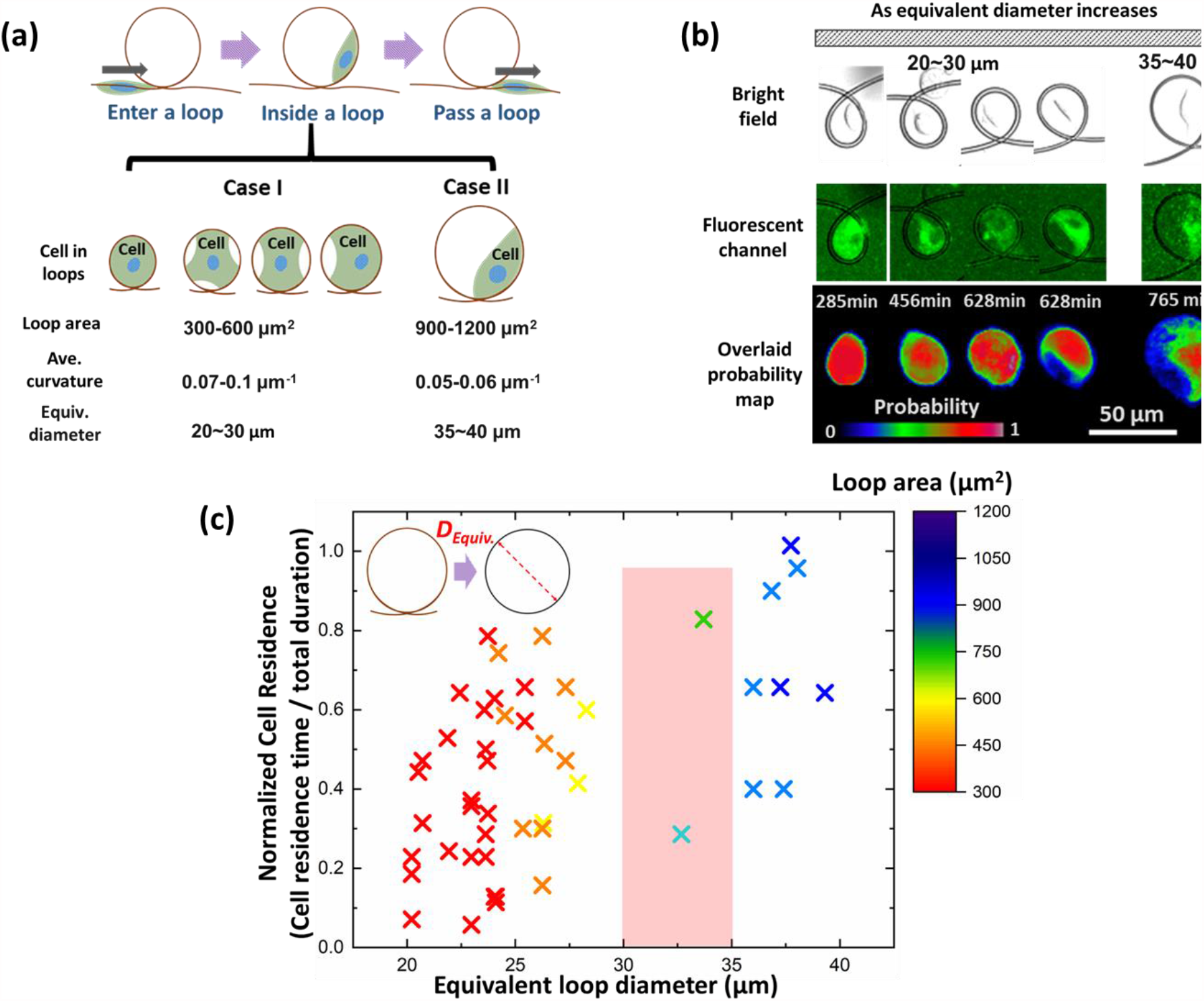
Analysis of single cell migrations within loop polystyrene microfibre patterns. (a) Schematic illustrations demonstrated distinct sizes of fibre loops, and the corresponding cell behaviours inside loops. (a) a sketch showing different stages of a MDA-MB-231 single cell migrating across a fibre loop, as well as distinct sizes of fibre loops, and the corresponding cell features inside loops; (b) Fluoresenct microscopic images and cumulative probability maps demonstrated distinct loop sizes affect single cell behaviours inside, scale bar=50 μm; (c) Scattered distribution between the normalized cell residence and the fibre loop size. Equivalent loop diameter is considered as the feature length; the 30-35 μm range is considered as the threshold feature length that indicate different cell behaviours.

## 3. Conclusion

For all the global microfiber patterns studied in this work, MDA-MB-231 cells were able to proceed fibre guided migrations, meanwhile adapting their morphologies to fit in local topographies as well as fibre global patterns. As shown in **Figure 7**(a), results above point to the threshold feature length for aligned, wavy, loop and grid fibre patterns are all at the range of 30-35 μm for which the MDA-MB-231 cells would switch its migration behaviour. It is intuitive to reason that this critical pattern feature size would be related to intrinsic cell shape features. Therefore, we examined the minor axis length and the major axis length of the cells in all experiments, as plotted in **Figure 7**(b)-(c). It is shown that the upper limit of the cell minor axis length, and the lower limit of cell major axis length, are both in the range of 30-35 μm. The ‘coincidence’ of these two values led us to postulate the following: A cell can have different movements to accommodate fibre patterns during migration; different fibre patterns will determine the motion as well as the shape of the cell. However, the chances for a cell to alter its pre-existing fibre migration path depend on its ability to stretch sideways and reach the adjacent fibre, and thus could be reflected by the upper limit of its minor axis. For MDA-MB-231 cells which attached to an existing fibre, the maximum distance that they can reach and form stabilised adhesion on another fibre, over the preference of existing fibre alone, is in the range of 30-35 μm.

**Figure 7.**
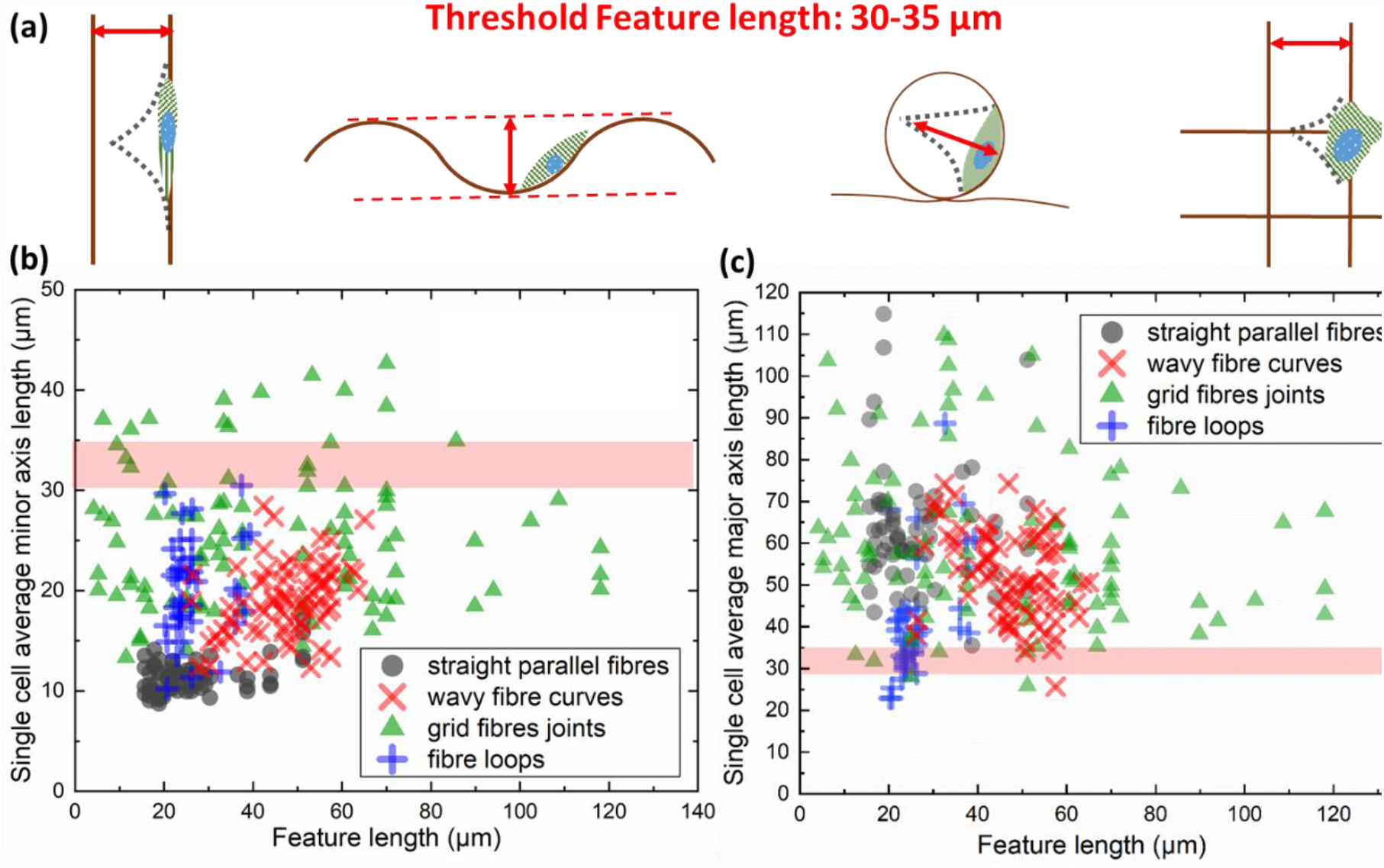
Feature length reflected the capability for MDA-MB-231 cells to stretch and adapt to the fibril environment. (a) Schematic diagrams showing the at different fibre patterns, the threshold feature length are all at the range of 30-35 μm; (b)-(c) Scattered plots of the average minor and major axis length for all single cells in different fibre patterns, revealing that the upper limit of the cells’ minor axis length, and the lower limit of cells’ major axis length, along with marking for the threshold feature length of 30-35 μm.

To conclude, we report the systematic analysis of MDA-MB-231 cell migration dynamics along polystyrene fibres fabricated using low-voltage continuous electrospinning. The fabrication of fibre networks with straight, curly and grid patterns provided well-defined and repeatable geometric environments to study how cells sense and perform interactions with such fibre morphologies. With such microfiber-deposited non-passivated substrates, cells were able to dynamically adapt their shapes in response to the adhered fibre, as well as to the neighbouring patterns; such adaptation reflect their ability to shuttle/transverse between fibre tracks. Key morphological features such as the variation of cell minor axis were identified for understanding cell migration in fibril matrices, as well as designing future ECM-mimic fibril models for cell migration studies.

## Experimental sections

### Coating glass slides

The glass slides (diameter: 22 mm) used for depositing polystyrene fibres were coated using a spincoater (Model WS-650MZ-23MPPB). For each glass slide, 70 μl 2 wt% polystyrene (Mw ∼ 280.000 kDA, Sigma-Aldrich) in toluene (anhydrous 99.8%, Sigma-Aldrich) solution was initially added at the centre. The glass slides were spun for 10 seconds at 500 rpm, followed by 50 seconds at 1000 rpm. Coated glass slides were then dried at reduced pressure for 24 hours, following which they could be stored at room temperature for several weeks.

### Fibre printing

Low-voltage continuous electrospinning patterning technology (LEP) was used to print polystyrene fibres on coated glass slides. The setup consists of a supporting aluminium plate and silicon plate under the glass slide, aluminium initiators at both sides of the glass slide, and a power source and syringe pump. Coated glass slides were placed between two aluminium initiators on a supporting aluminium plate for fibre deposition. The printing stage was heated up to 33.5°C to ensure continuous jetting during printing. After the stage had settled at this temperature, a 1 ml syringe with a No. 25 needle was filled with a 25 wt% polystyrene in dimethylformamide (Sigma-Aldrich) solution for fibre printing, and a flow rate of 160 μl/h was set. During the printing process, the stage was initially lifted slowly until it had reached the needle of the syringe. The positive and negative poles of the voltage output device were connected to the needle and the stage respectively. Polystyrene fibres were directly written onto the coated substrate under an applied voltage ranging from 0.3 kV to 0.7 kV. The global fibre pattern shape can also be affected by the voltage level applied. A voltage of of 0.5 kV marked the transition between straight and curly fibres for 25 wt% PS/DMF. To fabricate straight aligned fibre patterns, the working voltage was set at 0.3 kV. To fabricate curly fibres, working voltage was further increased to 0.7k V. To fabricate grid fibre patterns, a first layer of straight aligned fibres were initially deposited, followed by rotation of the substrate by 90 degrees to deposit a second layer of aligned fibres.

### Fibre pre-treatment

A series of pre-treatments were followed to prepare deposited fibres for cell adhesion and migration experiments. First, silicon rubber compound (RS 555-588) was used to fix the printed glass slide on a petri dish (diameter: 35 mm). This petri dish had a hole cut in the centre, using a machine in advance, and the glass slide was fixed right above this hole. After fixing, plasma treatment was applied with a flow rate of 25 m^3^/s for 45 seconds to provide polystyrene fibres with hydrophilic surface. Afterwards, the petri dish was sterilized under UV light for 30 minutes, following which the fibres were immersed in 4 ml PBS for 24 hours in the hood. Finally, PBS was replaced with 3 ml cell culture medium and the fibres were immersed for 48 hours in a standard cell incubator. During this step, the immersing culture medium was changed every 24 hours.

### Cell culture

GFP-labeled MDA-MB-231 human breast cancer cells were cultured in Dulbecco’s Modified Eagle’s Medium (DMEM, Invitrogen) supplemented with 1% Lglutamine (Invitrogen), 10% fetal bovine serum (FBS), 1% penicillin and 100 μg/ml streptomycin (Sigma Aldrich). To reduce fluorescence quenching during culture, aluminium foil was used to wrap the culture flasks. The culture medium was changed every 2 days, and cell passage was performed when 80% of the flask was covered by cells.

### Live cell imaging

The cell Z-stack imaging, live/dead assays and time-lapse experiments were performed using an environmentally controlled chamber joined to a Leica confocal microscope (Leica TCS SP5). The heater was set to 37°C to heat up the stage top incubator, and sterilized water and CO_2_ were used to provide a suitable moisture and pH inside the box-type incubator. During imaging, excitation was set at 480 nm and detection was set at around 500 nm to observe the GFP signal, and the bright field signal was also collected. A 20X objective lens was used, and up to 4 views were selected at one time. Cells were seeded on fabricated fibre patterns. An estimated period of 2-3 hours was given in advance before the imaging initiated, in order for most of the cells to attach to fibers; and their migration dynamics were captured every 10.5 minutes in the time-lapse microscopy for a continuous 12-hour time period.

### Post-Processing and Statistical Analysis

For the image analysis, an image analysis program written in-house was used to pre-process the live-cell images and to create a database to classify the microenvironment-based interaction automatically. The program incorporates a code to analyse the z-stack images, and a number of open-source applications, including ImageJ plug-ins and CellProfiler pipelines. Briefly speaking, as illustrated in **Figure S1**, the GFP intensity of the z-stack imaging acquired during the time-lapse experiments was projected into one x-y plane at each time point; thus the projected cell area, that is the 2D projection of the 3D cell shape, was obtained at each time point. The Manual Tracking plugin of ImageJ, applied to the projected GFP sequence, was used to track the position of the cancer cell nuclei over time. Cells that were long-lasting in the acquisition view, kept moving along single a fibre and were distanced from other cells, were generally preferred for tracking.

Home-written Matlab codes were used for most analysis of cell migration. Additionally, Matlab code provided by Wu et al.^36^ was used for persistent random walk model simulation.

The image analysis software CellProfiler was applied to perform the segmentation of the cancer cell projected area and to extract the cell shape features in the different microenvironments at each time point. The cancer cell coordinates were corrected to take into account the fluctuations of the microscope stage that occurred during experiments. The z coordinate of the cancer cells was calculated by finding the z level of the maximum GFP within the projected cell area.

## Supplementary Materials

**Figure S1.**
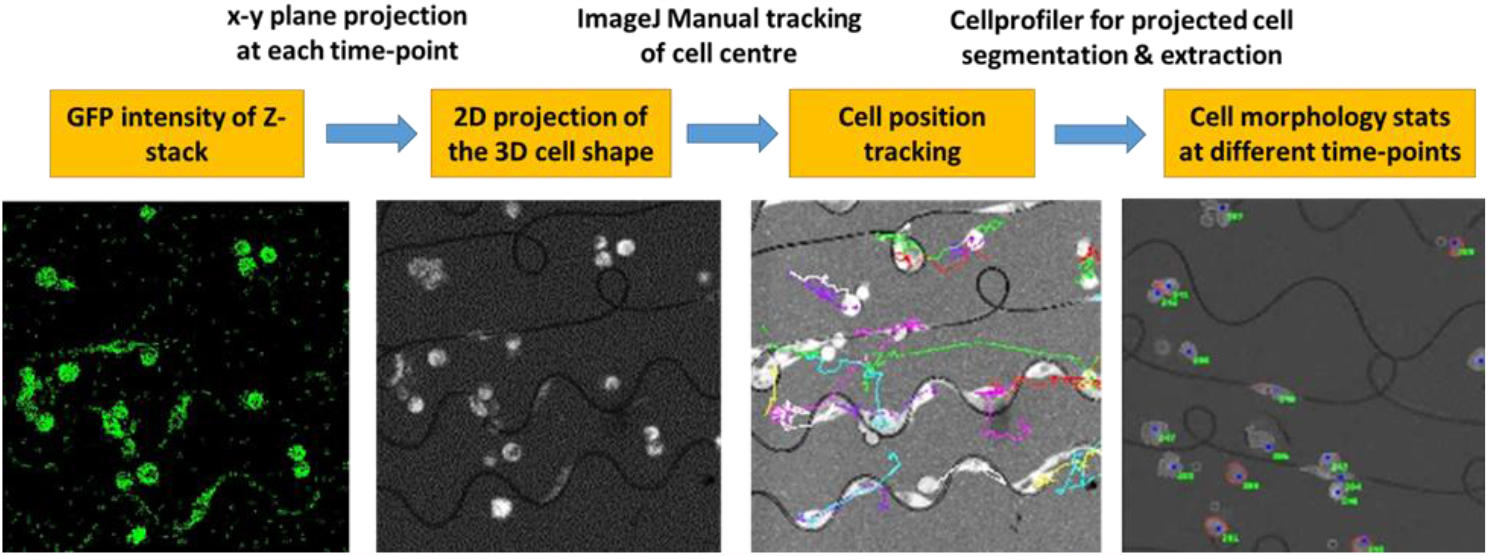
Flow chart demonstrating the procedure of the image processing steps based on the live-cell microscopy. The GFP intensity of the z-stack cell imaging acquired during the time-lapse experiments was projected into one x-y plane at each time point; thus the projected cell area, was obtained at each time point. ImageJ was used to track the position of the cancer cell nuclei over time, and CellProfiler was applied to perform the segmentation of the cancer cell projected area and to extract the cell morphological features at each time point.

**Figure S2.**
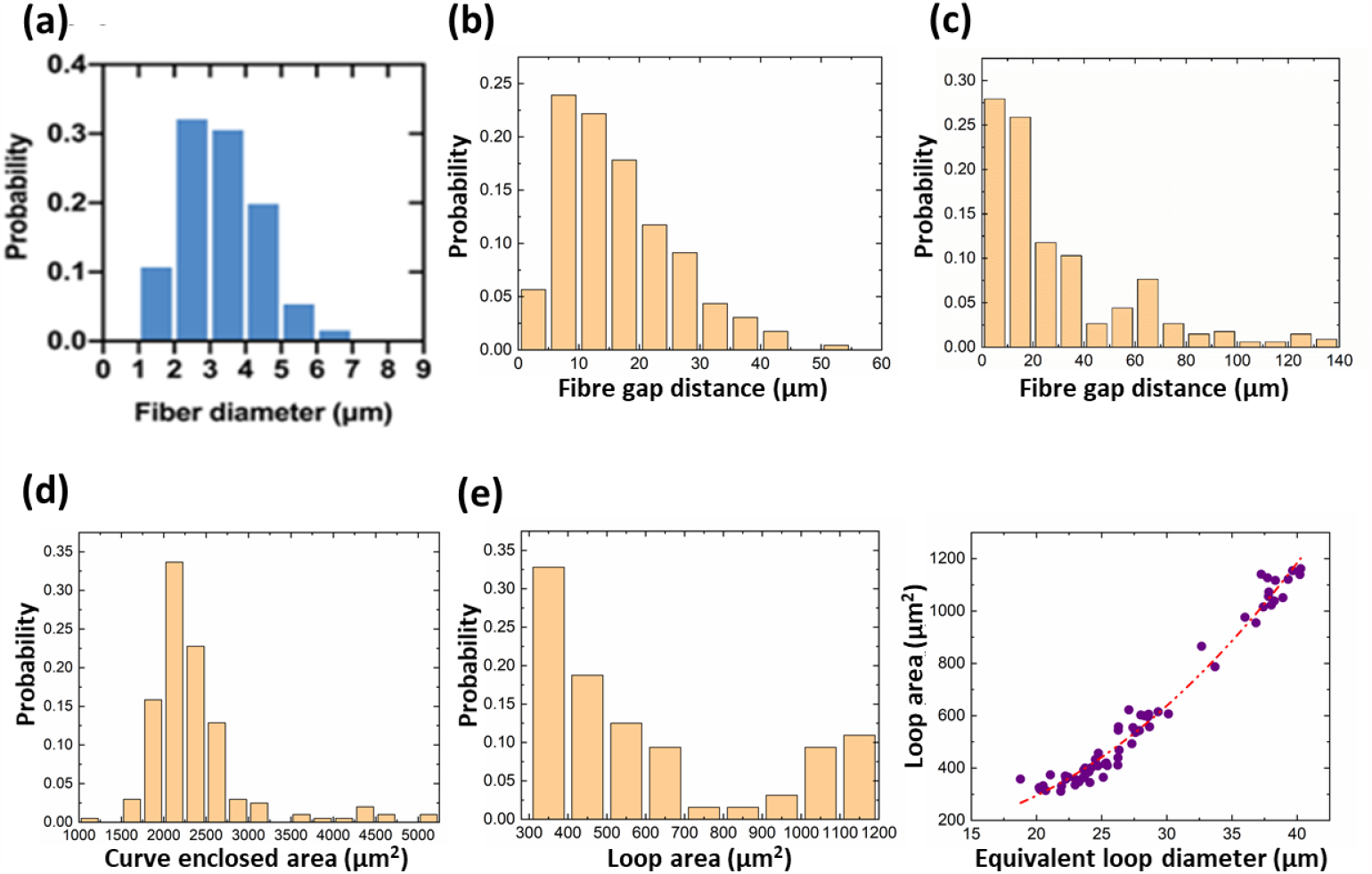
Set of diagrams demonstrating topographical characteristics of fabricated fibre patterns. (a) The distribution of fibre diameters for the straight fibres, mean S.D 3.3 ± 1.1μm; (b) For the straight aligned fibres, the probability distribution of inter-fibre distances, mean S.D 16.4 ± 9.5 μm; (c) For the grid fibre networks, the probability distribution of inter-fibre distances; (d) For the semilunar wavy fibre curves, the probability distribution of curves’ enclosed areas; (e) For fibre loops, the probability distribution of loop areas, as well as the correlation between derived loop average diameter and measured loop areas.

